# Mechanisms of life cycle simplification in African trypanosomes

**DOI:** 10.1101/2024.07.12.603250

**Authors:** Guy R Oldrieve, Frank Venter, Mathieu Cayla, Mylene Verney, Laurent Hébert, Manon Geerts, Nick Van Reet, Keith R Matthews

**Affiliations:** Institute for Immunology and Infection Research, School of Biological Sciences, University of Edinburgh, Edinburgh EH9 3FL, UK; York Biomedical Research Institute and Department of Biology, University of York, York; Unité Physiopathologie et Epidémiologie des Maladies Equines (PhEED), Laboratoire de Santé Animale, Site de Normandie, Agence Nationale de Sécurité Sanitaire de l’Alimentation, de l’Environnement et du Travail (ANSES), 1180 route de l’église, 14430 Goustranville, France; Department of Microbiology, Immunology and Transplantation, Rega Institute for Medical Research, Katholieke Universiteit Leuven, Leuven, Belgium; Department of Biomedical Sciences, Institute of Tropical Medicine, Antwerp, Belgium

## Abstract

African trypanosomes are important parasites in sub-Saharan Africa that undergo a quorum-sensing dependent development to morphologically stumpy forms in mammalian hosts to favour disease transmission by tsetse flies^1^. However, some trypanosome lineages have simplified their lifecycle by escaping dependence on tsetse allowing an expanded geographical range, with direct transmission between hosts achieved via biting flies and vampire bats (*Trypanosoma brucei evansi*, causing the disease ‘surra’) or through sexual transmission (*Trypanosoma brucei equiperdum*, causing ‘dourine’). Concomitantly, stumpy formation is reduced or lost, and the isolates are described as monomorphic, with infections spread widely in Africa, Asia, South America and parts of Europe. Here, using genomic analysis of distinct field isolates, we identified and functionally confirmed molecular changes that accompany the loss of the stumpy transmission stage in monomorphic clades. Further, using laboratory selection for resistance to the parasite’s quorum-sensing signal, we identified reversible steps in the initial development of monomorphism. This study identifies a trajectory of events that simplify the life cycle in emergent and established monomorphic trypanosomes, with impact on disease spread, vector control strategies, geographical range and virulence.

## INTRODUCTION

*Trypanosoma brucei* subspecies are the causative agents of human African trypanosomiasis (HAT) and Animal African trypanosomiasis (AAT). These parasites have a dixenous life cycle, entailing transmission between the mammalian host and the tsetse fly vector, this complex life history having evolved independently multiple times from monoxenous trypanosomatid ancestors ^2^. Dixenous *T. brucei* are pleomorphic, involving a developmental transition from a proliferative ‘long slender’ form to a cell-cycle arrested ‘short stumpy’ form, which is preadapted for uptake by the tsetse fly ^3,4^. The transition from slender to stumpy forms involves a density-dependent process where parasite-released peptidases hydrolyse extracellular proteins to produce oligopeptides as a quorum sensing signal ^5,6^. These are received via TbGPR89, and initiate a signalling cascade which induces developmental progression ^7,8^ in a process that can be recapitulated in vitro using oligopeptide-rich broth ^9^.

The complex lifecycle of *T. brucei* restricts these parasites to the geographic range of the tsetse fly within Africa. However, some clades have simplified their life cycle by foregoing development in their arthropod vector, allowing them to escape the tsetse belt, causing disease in Asia, South America, and Europe. These have been historically described as separate species, namely *T. evansi* and *T. equiperdum* ^10^, defined by their morphology, host species and disease presentation. *T. evansi* was described as the causative agent of the disease surra, exploiting mechanical transmission via biting flies, whereas *T. equiperdum* was described as the causative agent of dourine, being sexually transmitted between equids ^11^. Without tsetse transmission, each is characterised by their reduced production of arrested stumpy forms and so are described as monomorphic. Having dispensed with the tsetse fly vector and the associated metabolic needs at that stage of the life cycle, monomorphic *T. brucei* frequently exhibit a reduced or absent mitochondrial genome, the kinetoplast (kDNA), as a secondary consequence ^12^. Since *T. brucei* sexual reproduction occurs in the tsetse fly salivary gland ^13,14^, a further consequence of the simplified life cycle of monomorphs is that they are obligately asexual, the parasites being constitutively diploid in their mammalian host. Although this transition facilitates the expansion of monomorphic *T. brucei* into new ecological niches, in the long term obligate asexual reproduction prevents the organism from purging deleterious mutations whilst also preventing the maintenance of population-wide genetic diversity, hindering adaptation ^15,16^. Recent analyses revealed monomorphic *T. brucei* clades are polyphyletic, with each clade separated by pleomorphic isolates ^17–20^, and are currently most accurately described as ‘types’; *T. b. equiperdum* type BoTat, *T. b. equiperdum* type OVI, *T. b. evansi* type A and *T. b. evansi* type B ^19,20^. Although mechanisms to adapt to kDNA loss have been described ^21^, the molecular events which originate a simplified lifecycle are unknown.

## RESULTS

We reasoned that the occurrence of multiple naturally occurring monomorphic clades, alongside the ability to generate new monomorphic isolates by selection against stumpy formation in vitro, could provide insight into the mechanisms underlying simplification of the parasite lifecycle and the evolution of monomorphism. Therefore, 83 *T. brucei* isolates from diverse hosts such as humans, cattle, equids, camel and tsetse (Supplementary data S1), including 37 monomorphic isolates, were analysed for their phylogenetic relationships. The resulting phylogeny corroborates earlier evidence for at least 4 independent origins of monomorphic *T. brucei* (Fig. 1a) ^19,20^. As previously noted, the isolate IVM-t1, which was isolated from the genital mucosa of a horse and typed as *T. equiperdum* ^22^, groups with *T. b. evansi* type B isolates but shares a relatively distant relationship to other isolates designated in that clade, suggesting that IVM-t1 could represent a 5th independent monomorphic clade ^20^ (Fig. 1a).

**Fig. 1:**
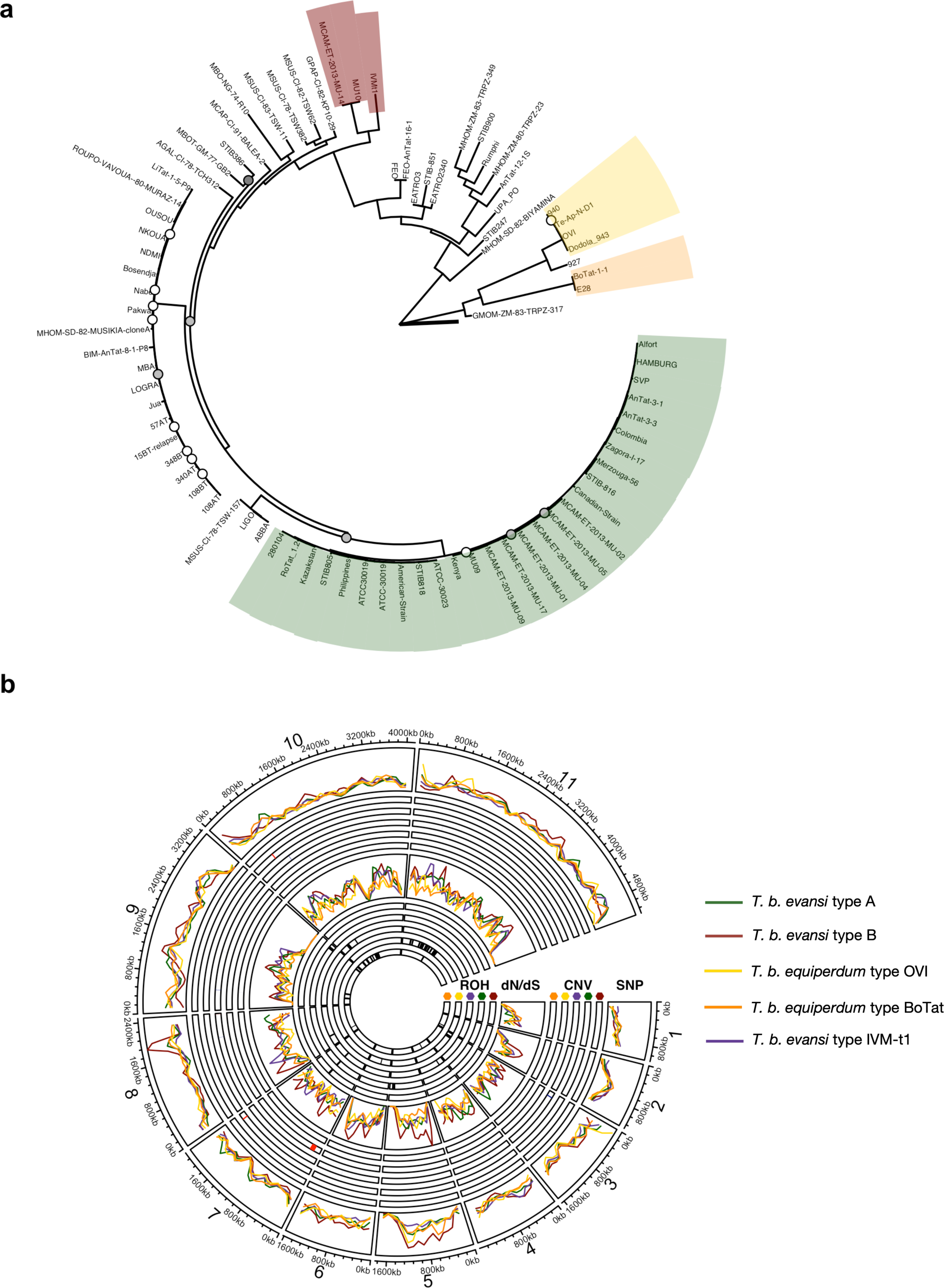
Monomorphic *T. brucei* are polyphyletic and form at least four independent clades, each displaying clade specific features at the genome scale. **(A)** A centrally rooted phylogenetic tree was created with 368,730 SNPs. Bootstrap confidence below 100 are reported by circles; white: 75-100, grey: 50-75 and black: 25-50. The tree scale, highlighted by the bold line at the tree root, represents 0.05 substitutions per site. Monomorphic isolates form at least four distinct clades, *T. b. equiperdum* type OVI, *T. b. equiperdum* type BoTat, *T. b. evansi* type A and *T. b. evansi* type B. IVM-t1, an isolate originating from the genital mucosa of a horse, groups with *T. b. evansi* type B. Pleomorphic clades are not highlighted. A full list of the isolates used in this tree is found in Supplementary Data S1. **(B)** The density of clade-specific mutations, location of copy number variation (CNV) highlighted in red (increase in copy number) or blue (decrease in copy number), density of genes with a positive dN/dS ratio (dN/dS) and locations of runs of homozygosity (ROH) highlighted in black. Each statistic is calculated across the 11 megabase chromosomes for each monomorphic clade. CNV (outer circles) and ROH (inner circles) are split into tracks for each clade ordered out to in; *T. b. evansi* type A (green), *T. b. evansi* type B (red), *T. b. evansi* type IVM-t1 (purple), *T. b. equiperdum* type OVI (yellow) and *T. b. equiperdum* type BoTat (orange).

### Independent monomorphic clades display unique hallmarks of transition to an obligate asexual lifecycle

At the chromosome level, we did not observe enrichment in the normalised density of mutations between clades, however, peaks and troughs of mutation density are found in smaller regions, such as *T. b. evansi* type B on chromosome 8 (Fig. 1b – SNP track). Distinct from *Leishmania* spp. where remarkable copy number variation (CNV) plasticity provides a mechanism to rapidly respond to environmental stimuli ^23^, we did not identify clade specific CNV in monomorphic *T. brucei* at the chromosome level (Fig. 1b – CNV track), corroborating previous CNV studies in *T. brucei* subspecies ^24^. However, *T. b. equiperdum* type OVI does display smaller CNV, the largest being a duplicated region of Chromosome 7. This region spans 46 genes, which are significantly enriched for gene ontology (GO) molecular functions; ‘glutathione peroxidase activity’, ‘peroxidase activity’, ‘oxidoreductase activity acting on peroxide as acceptor’ and ‘antioxidant activity’ (Fig. S1a).

Being obligately asexual and so unable to undergo recombination, we sought to identify evidence of the accumulation of deleterious mutations in monomorphic lineages by calculating the diversity of nonsynonymous to synonymous mutations (dN/dS). Although we did not find broad enrichment of genes with an accumulation of deleterious mutations at the chromosome level (Fig.1b – dN/dS track), each clade displayed >1,000 genes which had a positive selection (dN/dS >1), highlighting an accumulation of non-synonymous mutations in a monomorphic clade but neutral (dN/dS = 1) or purifying (dN/dS <1) selection in the pleomorphic background (Supplementary Data S3). Mitochondrial GO components were common amongst these sets of genes for all clades aside from *T. b. evansi* type B (Fig. S1b-f). *T. b. evansi* type B GO enrichment for mitochondrial membrane components fell below our significance threshold but was significantly enriched for membrane components (Fig. S1c). Only 55 genes (Supplementary Data S2) were under positive selection in all monomorphic clades, but these were not significantly enriched for GO components, broadly suggesting that monomorphism is associated with an accumulation of mutations in mitochondrial components, but that specific components are unique to each clade. Focusing on groups of known genes associated with the QS pathway or those with enriched expression in the insect stage of the parasite’s life cycle, QS genes have an increased dN/dS ratio in *T. b. equiperdum* type OVI and *T. b. equiperdum* type BoTat compared to essential genes (Fig. S1g).

Finally, we examined runs of homozygosity (ROH) which can highlight regions of the genome that have undergone gene conversion as a mechanism for asexual organisms to rid themselves of deleterious mutations in the absence of sexual recombination ^16^. This identified clade specific ROH on chromosomes 10 and 11 in *T. b. equiperdum* type OVI and *T. b. equiperdum* BoTat, respectively (Fig. 1b, - ROH track). The ROH in *T. b. equiperdum* type BoTat covers 1,004 genes but is not enriched for a particular GO term, whereas the 875 genes covered by the *T. b. equiperdum* type OVI ROH was enriched for GO components ‘transcriptionally silent chromatin’, ‘replication fork’, and ‘non-membrane bound organelles’ (Fig. S1h).

### Identification of molecular changes that reduce stumpy formation

To identify changes likely to contribute to the life cycle simplification of the monomorphic lineages, clade-specific mutations were experimentally targeted. These were selected based on (i) their presence in a monomorphic lineage but not in pleomorphic isolates, (ii) association with QS, and/or (iii) with a positive dN/dS ratio. Of these, an exemplar subset of genes were functionally characterised via a pipeline involving locus-specific replacement of both alleles in developmentally competent *T. brucei* AnTat1.1 using CRISPR-Cas9, and add-back replacement, with each step mirrored in parallel using wild-type allelic replacements to control for transfection and culture-associated loss of pleomorphism. For each cell line, developmental capacity was assessed using growth in vitro in the presence or absence of brain-heart infusion (BHI) broth as a source of QS oligopeptides ^5,25^, with cell cycle development and expression of the stumpy-specific marker protein PAD1 ^26^ monitored at the individual cell level. In combination these define stumpy formation, which is reduced or lost in monomorphic lineages exposed to BHI (Fig. S2) ^27^.

Of genes with a known involvement in QS signalling, replacement of pleomorphic alleles of Tb927.11.6600 (Hyp1), Tb927.8.1530 (TbGPR89) and Tb927.4.3650 (PP1) with monomorphic sequences did not alter developmental competence (Fig. S3b-d), although there was slow growth in parasites expressing the *T. b. equiperdum* type BoTat TbGPR89 variant (Fig. S3b). In contrast, a developmental phenotype was observed with some monomorphic clade specific Tb927.2.4020 alleles (Fig. 2a and S3a). This gene encodes the NEDD8-activating enzyme E1 (APPBP1), which along with UBA3, binds NEDD8 and activates protein neddylation, a highly conserved eukaryotic posttranslational modification pathway required for diverse cellular processes ^28–32^. Monomorphic clade-specific non-synonymous mutations in APPBP1 are found in *T. b. evansi* type A (n=4), *T. b. evansi* type IVM-t1 (n=5) and *T. b. equiperdum* type BoTat (n=4), although the structural consequences of each were predicted by AlphaFold (*26*) to be minimal (Fig S4a,b). In standard media (HMI-9), cells expressing APPBP1 sequences from the monomorphic lineages *T. b. evansi* type A and *T. b. equiperdum* type BoTat, control pleomorphic (TREU927/4) or pleomorphic add-back displayed no significant difference in growth over 72 hours, although there was growth delay in the cell line expressing the *T. b. evansi* type IVM-t1 monomorphic APPBP1 sequence at 48 hours (Fig.2a and Fig. S3a). In the presence of the QS oligopeptide signal, BHI, parasites expressing the *T. b. equiperdum* type BoTat and *T. b. evansi* type A sequences arrested growth equivalently to those expressing the pleomorphic *T. b. brucei* alleles (Fig. S3a). In contrast, cells expressing the *T. b. evansi* type IVM-t1 APPBP1 alleles continued to grow when exposed to BHI (Fig. 2a), indicative of reduced sensitivity to the QS signal. This correlated with a higher proportion of cells with 2 kinetoplasts and one or two nuclei (indicative of cells in active replication) and the absence of expression of the stumpy stage-specific protein, PAD1, in contrast to cells expressing the *T. b. brucei* sequence (Fig. 2b). Confirming the phenotype was dependent upon the monomorphic APPBP1 sequence, add-back of the pleomorphic *T. b. brucei* sequence to replace the *T. b. evansi* type IVM-t1 sequence in the cell line restored growth arrest and PAD1 expression once exposed to BHI (Fig.2a-b).

**Fig. 2:**
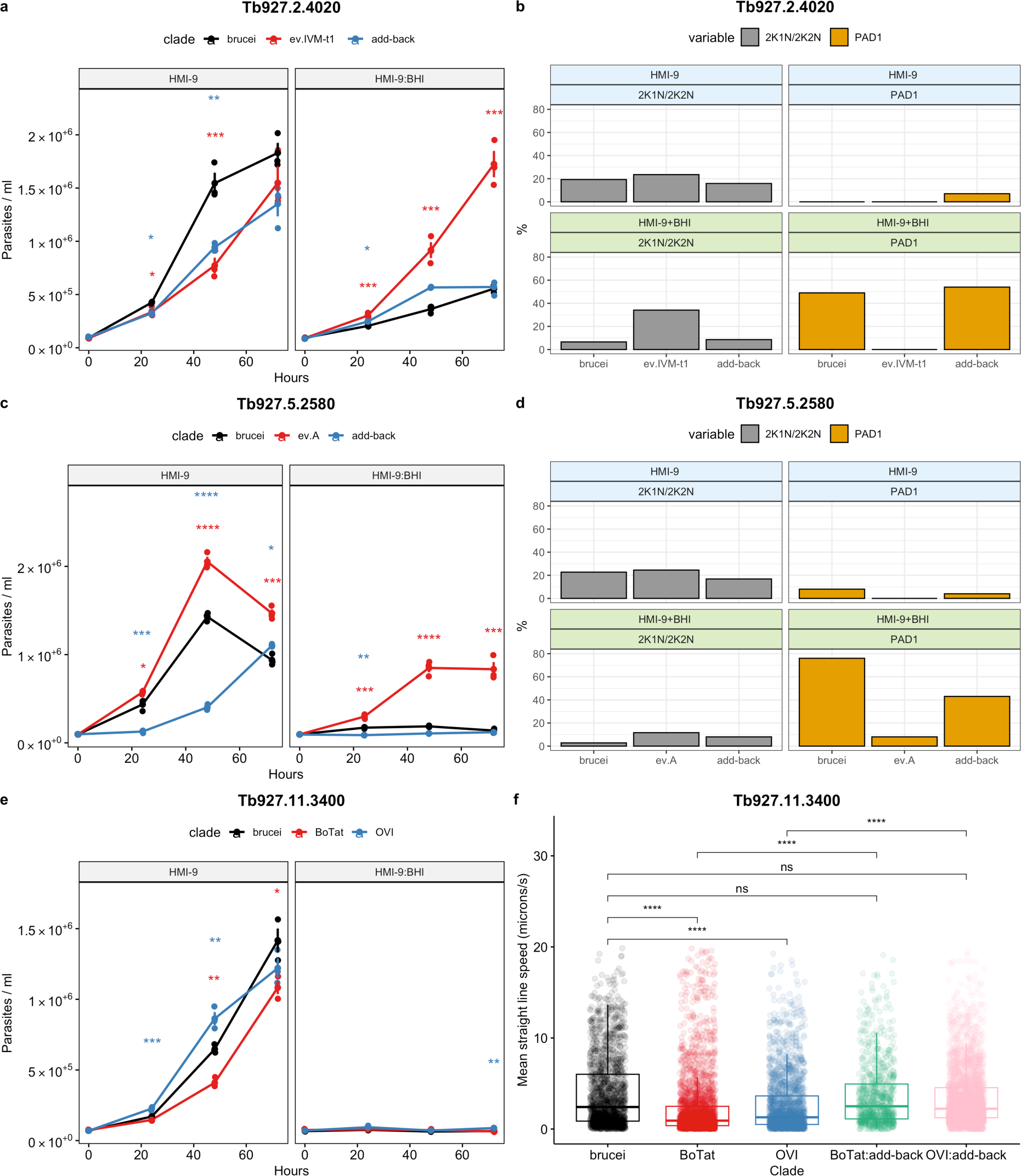
Monomorphic *T. brucei* clades display mutations in discrete genes which hinder pleomorphic phenotypes. **(A, C & E)** Growth and **(B & D)** percentage of the population replicating and PAD1+% from pleomorphic *T. brucei* EATRO 1125 AnTat1.1 J1339 subjected to replacement of endogenous gene targets **(A-B)** Tb927.2.4020 APPBP1, **(C-D)** Tb927.5.2580 – hypothetical protein and **(E-F)** Tb927.11.3400 - Flagellum attachment zone protein 41 (FAZ41). The replacement cell lines were grown in HMI-9, or HMI-9 supplemented with an in vitro mimic of the QS signal, oligopeptide broth BHI (15%). Significance at each timepoint and for each comparison is shown (brucei vs monomorph, *; brucei vs add-back, *). * (p<0.05); ** (p<0.01; *** (p<0.001); **** (p<0.0001). For the growth and IFA analysis, four flasks were grown, three for growth analysis and one to screen PAD1 and K/N counts at 48 hours. **(F)** The motility of the FAZ41 replacement cell lines was also quantified.

Interestingly, despite their contrasting phenotypic effects, the *T. b. evansi* type A and *T. b. evansi* type IVM-t1 APPBP1 sequences differ by only a single non-synonymous mutation, G224S (Fig. S3a). A cell line was generated that expressed APPBP1 with a sequence identical to pleomorphic *T. b. brucei* but with the single G224S mutation specific to *T. b. evansi* type IVM-t1. These cells arrested their growth upon exposure to BHI equivalently to those expressing the pleomorphic *T. b. brucei* sequence (Fig. S3e). Hence, the G224S mutation alone is not sufficient to reduce developmental competence but operates in the context of the other mutations in the *T. b. evansi* type IVM-t1 APPBP1 sequence.

Of those novel genes not previously linked to the QS response but identified based on their evidence for selection, Tb927.5.2580 has a positive dN/dS ratio in the monomorphic clade *T. b. evansi* type A. The sequence contains four clade-specific non-synonymous mutations compared to the genome reference strain TREU927/4 (Fig. S4g) and encodes a mitochondrially-located ‘MIX’-associated protein that cofractionates associated with the cytochrome c oxidase complex (complex IV) ^33^ which is expressed in procyclic forms but repressed in bloodstream forms ^34^. Interestingly, cells expressing the *T. b. evansi* type A Tb927.5.2580 sequence grew more rapidly in HMI-9 media compared to cells expressing the *T. b. brucei* sequence (Fig. 2c) and, in the presence of the QS signal, continued proliferation and displayed fewer PAD1+ cells contrasting with cells expressing the pleomorphic sequence (Fig. 2c-d). Supporting its contribution to development, homozygous add-back of the *T. b. brucei* sequence caused parasites to reduce their growth rate in HMI-9 media and restored arrest after exposure to BHI (Fig. 2c). This was accompanied by reduced 2K1N, 2K2N cells and increased PAD1 expression although not fully to wild type levels likely due to the number of transfection cycles (Fig. 2d).

Confirming these in vitro assays, analysis in vivo demonstrated that parasites with the *T. b evansi* type A Tb927.5.2580 sequence showed reduced arrest and stumpy formation compared to parasites expressing the pleomorphic alleles and re-introduction of the wild type allele reversed this (Fig. 3 & Fig. S5). In the *T. b. evansi* type A clade, Tb927.5.2580 contains one non-synonymous homozygous mutation which is unique, A149P. Expression of the *T. b. brucei* pleomorphic sequence with just the A149P mutation did not impact developmental competence (Fig. S3f). Hence, as for the G224S mutation in APPBP1, A149P appears to take effect in the context of other changes in the gene sequence rather than in isolation.

**Fig. 3:**
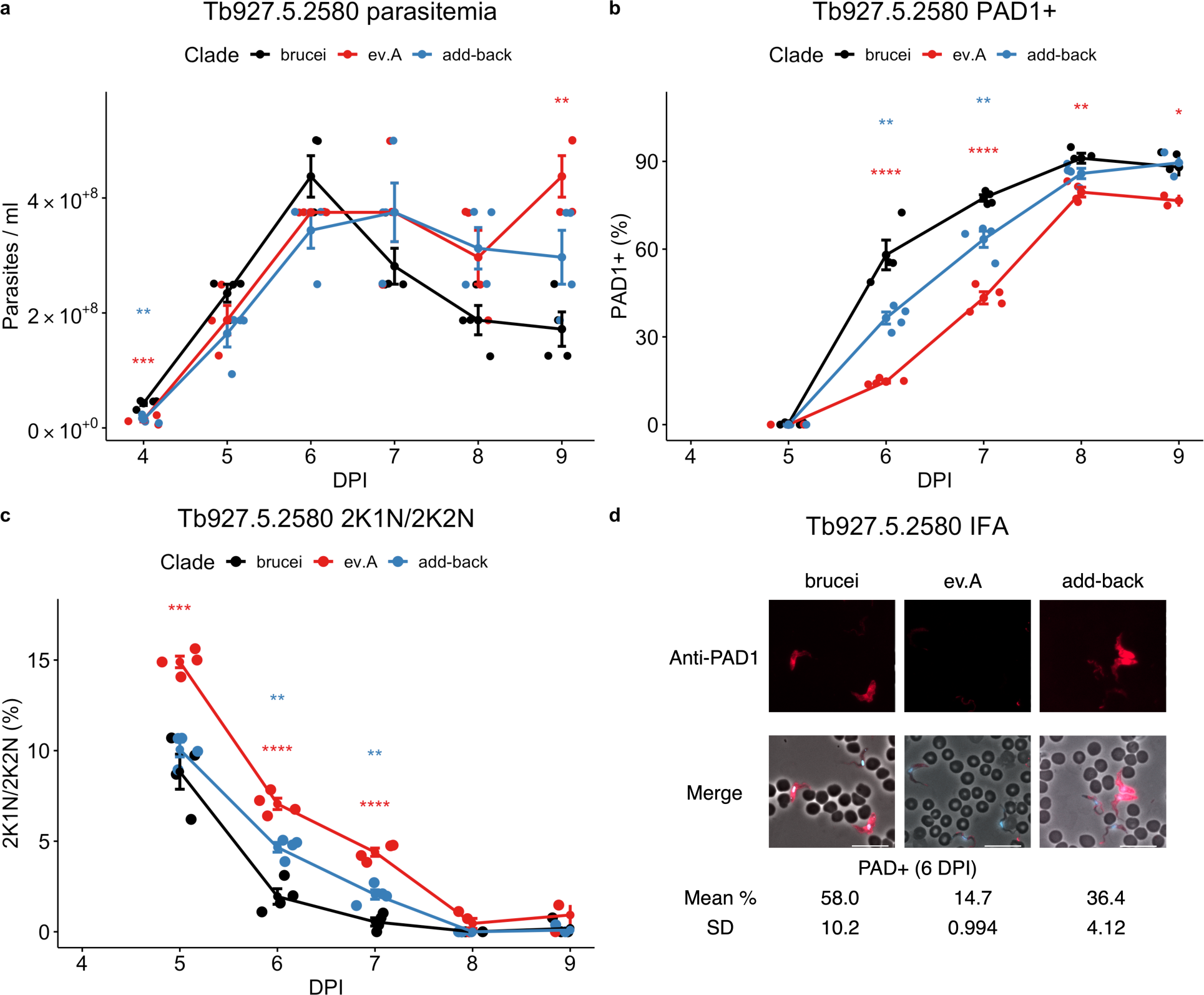
Pleomorphic *T. b. brucei* expressing the monomorphic *T. b. evansi* type A Tb927.5.2580 sequence delays developmental progression. **(A)** In vivo growth of pleomorphic *T. b. brucei* expressing the monomorphic *T. b. evansi* type A Tb927.5.2580 sequence or the pleomorphic *T. b. brucei* sequence, **(B)** cell cycle stage and **(C)** percentage of PAD1+ cells as assessed by immunofluorescence. Each cell line was used to infect four mice, represented by the dots at each time point. Error bars represent mean standard error. Significance at each timepoint and for each comparison is shown (brucei vs monomorph, *; brucei vs add-back, *). * (p<0.05); ** (p<0.01; *** (p<0.001); **** (p<0.0001). **(D)** Representative DAPI, PAD1 and merged images of each cell line at 6 DPI. Scale bar = 30µm.

Analysis of monomorphic-specific alleles that also displayed a developmental phenotype in a high throughout *T. brucei* phenotypic screen ^35^ highlighted Tb927.11.3400 which encodes FAZ41, a component of the trypanosome flagellum attachment zone (FAZ). *T. b. equiperdum* type OVI and *T. b. equiperdum* type BoTat share five identical non-synonymous mutations (Fig. S4l). Although each is believed to use sexual transmission, these isolates are of independent origin, phylogenetically separated by the pleomorphic TREU927/4 isolate (Fig. 1a). Replacement of FAZ41 with the *T. brucei brucei*, *T. b. equiperdum* type BoTat and *T. b. equiperdum* type OVI sequence did not alter developmental competence (Fig.2e). However, cells expressing either the *T. b. equiperdum* type OVI or *T. b. equiperdum* type BoTat sequence displayed significantly reduced motility in comparison to those expressing the *T. b. brucei* sequence. Add-back of the pleomorphic sequence restored motility to both initial replacement cell lines (Fig.2f; Supplementary movies 1-5). Given that monomorphic *T. b. evansi* (unspecified type) show altered motility as potential adaptation to the tissue environment ^36^, this highlights that our genetic comparisons can identify known phenotypic changes linked to monomorphism beyond developmental mechanisms. This was further corroborated by identification in our genome analyses of the F1 ATPase γ subunit (Tb927.10.180) mutations that permit kDNA loss previously identified in some monomorphic lineages ^21^. A full list of genes with a clade specific high-impact or moderate-impact mutation in a monomorphic clade along with a dN/dS ratio <=1 in pleomorphic isolates and >1 in any monomorphic clade is provided in Supplementary data 2, with predicted gene and protein alignments for all genes from all isolates accessible via GitHub (https://github.com/goldrieve/Mechanisms-of-life-cycle-simplification/blob/master/alignment_fasta/alignment.fasta.tar.gz).

### Gene regulators are downregulated as monomorphism is selected

The presence of multiple co-dependent and context-specific mutations in different genes and clades, which impact developmental competence, could act to lock parasites in the monomorphic state but are unlikely to be responsible for the original progression toward monomorphism. To explore how monomorphism might arise in the first instance, we used a laboratory selection protocol to isolate developmentally incompetent parasites de novo. Specifically, we independently selected monomorphic *T. brucei* cell lines via parallel serial passage of clonal lines in increasing concentrations of BHI (Fig. 4a), and then generated clones from the selected lineages after 30 passages. The reduced developmental competence of each selected line was then validated in vitro via exposure to BHI (Fig. 4a). In each case a reduced QS response was observed.

**Fig. 4:**
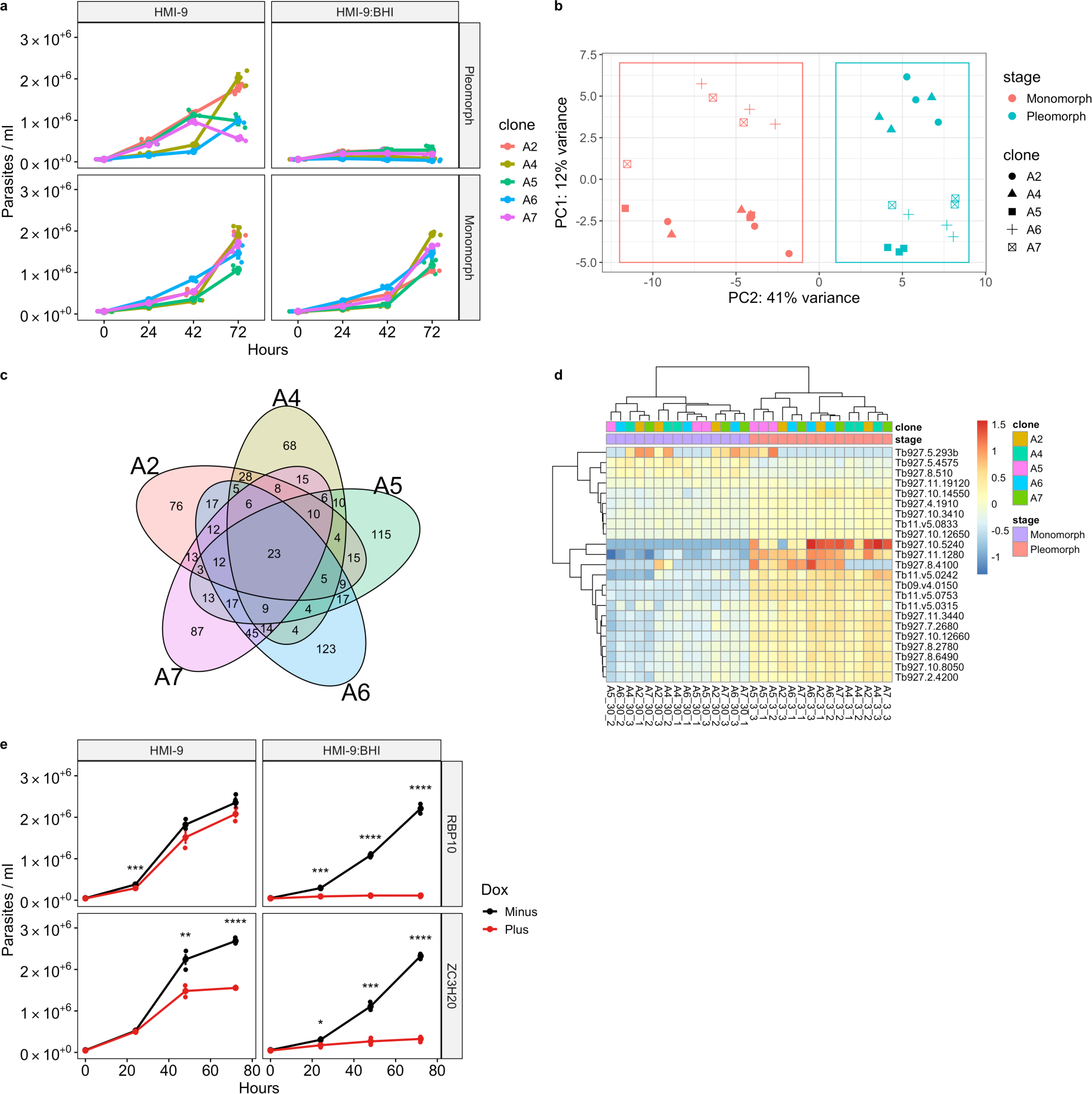
Selected monomorphs display reduced developmental progression, which is reversible by inducible expression of key gene regulators. **(A)** Five independent clones, derived by passage in increasing concentrations of BHI, display reduced developmental competence. The clones were grown in either HMI-9 or HMI-9 supplemented with 15% BHI; lines after passage 3 (pleomorph) or passage 30 are shown (monomorph). **(B)** PCA plot highlights the majority of variance in RNAseq libraries from the selection is explained by the selection from pleomorph to monomorph. **(C)** 23 differentially expressed genes are shared between each of the five clonal selections for monomorphism. **(D)** Heatmap of the 23 differentially expressed genes. **(E)** Growth of monomorphic clone A7 induced with doxycycline (Plus) or not (Minus) to express transgenic RBP10 and ZC3H20. Each cell line was grown in triplicate, represented by the dots at each time point. A triangle represents the mean for each time point. Error bars represent mean standard error.

Thereafter we analysed by the transcriptome profile of each clonal line before and after selection. This demonstrated that the pleomorphic progenitors and monomorphic descendants are clearly discriminated and clustered with respect to their developmental competence, indicating a common transcriptome trajectory among distinct selected lineages; overall, the transition from pleomorphism to monomorphism accounted for 41% of the transcriptome variance (Fig.4b). However, only 23 genes were significantly differentially expressed between all the selected clones (Fig. 4c, 4d). These genes were enriched for gene regulators, with GO terms including ‘regulation of gene expression’, ‘posttranscriptional regulation of gene expression’ as well as ‘regulation of macromolecule metabolic processes’ (Fig. S6d). Interestingly, the cohort of regulated transcripts included RBP10 (Tb927.8.2780), a known developmental regulator in trypanosomes ^37^ (Fig. S6b), which is itself predicted to bind ^38^ eight of the 23 commonly differentially expressed genes (Tb927.10.3410, Tb927.10.8050, Tb927.2.4200, Tb927.4.1910, Tb927.11.1280, Tb927.7.2680, Tb927.8.2780, Tb927.8.6490). Moreover, analysis of the expression of a manually curated list of quorum sensing (QS) genes (Fig. S6a) highlighted the key regulator ZC3H20 (Tb927.7.2660) ^9,39^, as well as PKA-R (Tb927.11.4610) and adenylosuccinate synthetase (Tb927.11.3650) ^7^ are differentially expressed during the selection for monomorphism (Fig. S6a).

We envisioned that the downregulation of RBP10 and its targets, alongside additional QS regulators such as ZC3H20, could initiate the first steps in the development of monomorphism, potentially reversibly in the first instance. To explore this, we re-expressed either RBP10 or ZC3H20 in a selected monomorphic line (A7) using a doxycycline-inducible expression system and assayed their responsiveness to the BHI QS signal. Fig.4e demonstrates that inducible expression of either ZC3H20 or RBP10 in the selected monomorphic clone A7 resulted in growth and cell cycle arrest of each in response to BHI (Fig. 4f). The expression of these individual RNA regulators in the selected line was not, however, sufficient in isolation to restore PAD1 expression (Fig. S6e).

## DISCUSSION

The generation of stumpy forms is maintained in *Trypanosoma brucei* populations to optimise transmission by tsetse flies ^40^. However monomorphic isolates have been able to simplify their life cycle by excluding the tsetse fly, allowing them to expand their geographical range via direct transmission between mammalian hosts. Here we have analysed the genomes of field isolates to identify molecular adaptations contributing to this simplified life cycle, identifying mutations in known regulators and previously uncharacterized QS components, as well as changes in genes affecting growth and motility. Moreover, multiple mutations within individual genes affecting stumpy formation were observed, which in APPBP1 and Tb927.5.2580 were phenotypically co-dependent. A similar phenotype was observed for PAG3, which was found to be disrupted in some monomorphic clades ^41^. As the deterioration of APPBP1, Tb927.5.2580 and PAG3 are clade specific, we suggest these mutations did not cause the monomorphic phenotype, but rather arose secondarily to the transition to monomorphism.

To explore the initiation of this phenotypic change, we performed laboratory selection to generate monomorphism de novo. In five independently selected clonal lineages this generated consistent changes in the transcriptome, with key posttranscriptional regulators showing altered expression, including two with established roles in life cycle progression (RBP10, ZCH20); previous selections for monomorphism have also identified changes in posttranscriptional regulators ^42,43^. Interestingly, we show that in selected monomorphs ectopic re-expression of the regulators allowed restoration of sensitivity to BHI, demonstrating some reversibility of the phenotypic change to monomorphism in these newly selected lineages. This resembles the sensitivity of *T. b. evansi* type A RoTat1.2 to BHI, where growth is inhibited without PAD1 expression (Fig. S2), perhaps indicating an early stage of monomorphism in that lineage.

In the field, we predict that monomorphism is initially reversible via changes in the expression of posttranscriptional regulators, with parasites (‘proto-monomorphs’) able to sustain transmission flexibility either through their tsetse vector or directly, potentially providing a selective advantage where tsetse transmission is challenging (Fig. 5). However, where tsetse become scarce through altered land use, tsetse control efforts, host migration or climate change ^44^, the proto-monomorphs continue to evolve as obligate asexual organisms. Mullers ratchet suggests that the organisms will be unable to purge these mutations through sexual recombination ^16^. Eventually, the accumulation of deleterious mutations in key QS genes, and the loss of their kDNA ^45^, ensures the loss of capacity for vectorial transmission and parasites become ‘locked’ as monomorphs. This has the short-term advantage of sustaining transmission in the absence of tsetse and expands the geographical range of the parasites into Asia, South America, and Europe.

**Fig. 5:**
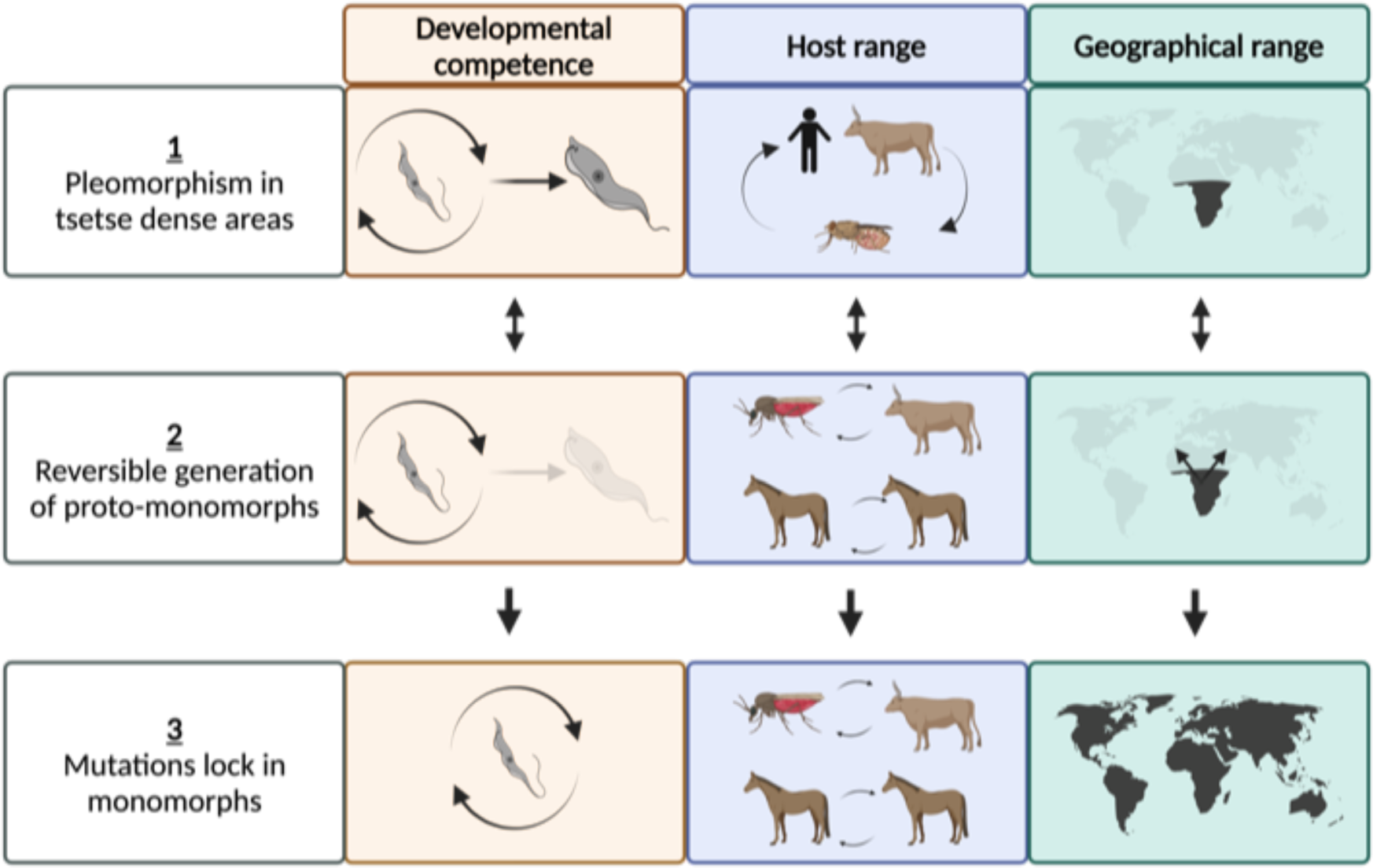
A model for life cycle simplification in *Trypanosoma brucei*. This proposes that (1) in tsetse dense areas, developmental competence is maintained, supporting the genetic diversity of the parasite population through sexual recombination in the tsetse vector. (2) If tsetse numbers fall, or infected hosts move from a tsetse endemic area, transmission flexibility can be favoured by downregulating developmental regulators, forming proto-monomorphs, promoting enhanced parasitaemia and so mechanical transmission. (3) With prolonged mechanical transmission, mutations accrue which would eventually lock the proto-monomorphs into an obligate asexual/ monomorphic lifecycle. Created with BioRender.com.

Whilst the accumulation of deleterious mutations would eventually doom the monomorphic lineages to extinction, our data suggests that new monomorphic variants may arise frequently and without detection, given their initial flexibility in transmission mode, and prior to the emergence of secondary characteristics such as kDNA loss. Given the drive toward enhanced virulence by adopting mechanical transmission ^46^, it will be important to detect emergent monomorphic lineages where tsetse population control strategies and climate change are modifying the range of the vector ^47^, particularly for zoonotic *Trypanosoma brucei rhodesiense* ^48^. The molecular events we have identified that underpin this process can also provide the necessary diagnostic tools ^11^ to detect and anticipate this threat.

## Supporting information

WT motility

OVI motility

OVI add back motility

BoTat motility

BoTat add back motility

## Methods

All code written for this study has been deposited on GitHub (https://github.com/goldrieve/Mechanisms-of-life-cycle-simplification). The workflow was implemented with SnakeMake ^49^.

### Parasite cell lines

Publicly available genome data were accessed from NCBI (SRA) ^50^, the Wellcome Sanger Institute or generated as part of this study. Isolates sequenced during this study were propagated as bloodstream-form populations in OF1 mice (Charles River, Belgium) and purified from infected mouse blood via DEAE ion exchange chromatography; we recognize that some isolates described as pleomorphic in this analysis may have undergone adaptation which potentially hindered their developmental competence as part of their DNA isolation steps and prior to genomic analysis. Purified trypanosomes were sedimented by centrifugation (3,000 × g, 10 min at 4◦C) and DNA was extracted by standard phenol-chloroform. The concentration of extracted DNA was determined using a Qubit 4 Fluorometer (Invitrogen by Thermo Fisher Scientific). Paired end 150 bp sequences were generated using the DNA nanoball sequencing technology (DNBSEQ™) at the Beijing Genomics Institute (BGI). Animal experimentation procedures were approved by the Ethics Committee for Animal Experimentation (ECD-ITG) at the Institute of Tropical Medicine in Antwerp, under proposal number DPU-2017.

### Variant calling

The quality of the raw reads was analysed with FastQC ^51^ and subjected to quality trimming with Trimmomatic ^52^. The trimmed reads were aligned to the *T. brucei* TREU927/4 V5 reference genome with bwa-mem ^53^. The reads were prepared for variant calling following the GATK4 best practices pipeline, which included marking duplicate reads ^54,55^. The variants were combined and filtered with stringent cut-offs, in keeping with GATK’s best practices pipeline and previous studies ^19,20^. The complete list of genomes analysed in this study, including their accession IDs, is summarised in Supplementary Data S1. Clade-specific mutations were assigned to annotated genes in the TRUE927/4 reference genome using snpEFF ^56^. Protein domains and putative phosphorylation sites were identified in target genes using InterProScan accessed through TriTrypDB (www.TritrypDB.org). snpSIFT was then used to identify mutations which were specific to a monomorphic clade using the case-control function ^56^.

### Phylogenetic analysis

The filtered variants, described above, were filtered again to retain SNPs where a genotype had been called in every sample, using VCFtools ^57^. A concatenated alignment of each variant was extracted using VCF-kit ^58^. IQ-TREE ^59^ was used to create a maximum-likelihood tree from homozygous variant sites. Within the IQ-TREE analysis, a best-fit substitution model was chosen by ModelFinder using models that included ascertainment bias correction (MFP+ASC) ^60^. ModelFinder identified TVM+F+ASC+R3 as the best fit and 1000 ultrafast bootstraps generated by UFBoot2 ^61^. The consensus trees were visualised and annotated with ggtree ^62^

### Genome feature summary

vcf2fasta.py (available here https://github.com/santiagosnchez/vcf2fasta) was used to generate a fasta file of each gene for each isolate based on TREU927/4 gene annotations. Pseudogenes and VSGs were removed from the analysis. Codeml ^63,64^ was then used to calculate the dN/dS ratio for each monomorphic clade, using the variants and phylogenetic tree created above. Only genes with a perfect alignment were included in the analysis, to avoid false positive results from misaligned sequences. The ctl file for each gene included: noisy = 3, verbose = 1, seqtype = 1, ndata = 1, icode = 0, cleandata = 0, model = 2, NSsites = 0, CodonFreq = 7, estFreq = 0, clock = 0, fix_omega = 0, omega = 0.5.

The dN/dS ratio was summarised for each clade and specific interest was taken for three groups of genes; (1) QS pathway genes, (2) genes used during the insect stage of the parasite’s lifecycle, specifically those which were significantly differentially expressed between parasites extracted from the midgut/proventriculus stage and proventriculus/salivary gland stage ^65^, (3) essential genes, defined as those with a significant reduction in every library of the RITseq high throughput phenotype study ^66^. The data was summarised in R.

Runs of homozygosity were identified with plink ^67^ using the following parameters: --homozyg-snp 20 --homozyg-kb 50 --homozyg-density 10 --homozyg-window-snp 10 --homozyg-window-missing 4 --homozyg-het 0 --maf 0.01 --homozyg-group --allow-extra-chr --family --write-cluster --cluster.

Dicey chop ^68^, bwa mem ^53^, samtools ^69^ and dicey map were used to create a mappability map for the TREU927/4 reference genome. CNVs were identified, merged and genotyped with delly ^70^ and outputs were merged again with bcftools ^71^. Delly was used to classify each CNV and summarised with bcftools.

The density of 17,820 clade-specific variants (17,820 variants were chosen as this represents the lowest number of variants found in any of the clades), the density of genes with a positive dN/dS ratio, locations of ROH locations of CNV were visualised for each monomorphic clade with circilize ^72^.

### Target prediction

Clade-specific variants described above were prioritised to create a target list of genes to validate their role in monomorphism. Initially, pseduogenes, VSGs and genes not on the megabase chromosomes 1-11 were removed. These genes were further filtered to create two target categories using the following criteria:

1. Genes which have a clade specific high-impact or moderate-impact mutation in a monomorphic clade in a known QS pathway gene ^5,7,8,73,74^. The gene must also display no high impact mutations in any pleomorphic isolates.
2. Genes which have a clade specific high-impact or moderate-impact mutation in a monomorphic clade along with a dN/dS ratio <=1 in pleomorphic isolates and >1 in any monomorphic clade. The gene must also have a smaller log fold change in D3, D6 and PF than in the DIF category whilst displaying a log fold change in the DIF category greater than −1.5 ^66^. The gene must also display no high impact mutations in any pleomorphic isolates.

After filtering, 38 genes from Category 1 and 540 genes from Category 2. The full lists can be found in Supplementary data S3. Six genes were taken forward for validation (four genes from Category 1 and two genes from Category 2.

### Protein structure prediction

Representative monomorphic mutant and pleomorphic control amino acid sequences were used to generate a protein structure model with ColabFold ^75^, which uses AlphaFold2 ^76^ and Alphafold2-multimer ^77^. The protein models were created using default settings and the best model, based on a predicted local-distance difference test (pLDDT), was aligned, and visualised with PyMOL ^78^.

### Synthesis of target replacement sequences

From our target list, we confirmed that all the target protein sequences were identical within the clade of interest and the sequence from a representative isolate from each clade was then chosen to synthesise a nucleotide sequence (Genewiz Gene Synthesis). As a control for each of these targets, the pleomorphic TREU927/4 sequence was also synthesised. To aid the downstream processing of the synthesised sequences, we added the enzyme sites HindIII and BamHI at the 5’ and 3’ terminus, respectively. HindIII and BamHI sites were identified within the coding sequence of Tb927.8.1530 and Tb927.2.4020, respectively. The sites were re-coded to remove the enzyme site whilst maintaining codon usage.

### Generation of plasmids required for CRISPR/ Cas9 transfection

The synthesised monomorphic sequences were generated in the pUC-GW-Kan plasmid (Genewiz) and transformed into XL1-competent cells. Plasmid preparation was performed using the GeneJET Plasmid Miniprep Kit (Thermo Fisher), following the manufacturer’s instructions. The plasmid was then digested using the HindIII and BamHI high-fidelity enzymes (NEB). The digested gene was excised from an agarose gel using the Monarch DNA Gel Extraction Kit (NEB). The gene was inserted into a pPOT plasmid using T4 DNA Ligase (NEB). Blasticidin (pPOT V6), hygromycin (pPOT V7), phleomycin (pPOT V7) and G418 (pPOT V7) resistance plasmids ^79,80^ using T4 DNA Ligase (NEB). Blasticidin, hygromycin, phleomycin and G418 resistance pPOT plasmids ^80^ were used at different stages throughout the following experiments, however, the same drug resistance plasmid was used for each set of experiments on a given gene. Hygromycin and Blasticidin were used to replace the wild-type alleles of every target gene apart from TbGPR89, for which we used Hygromycin and Phleomycin. Add-back of wild-type sequences to the initial replacement cell lines was performed using Phleomycin and G418. The following concentrations of each drug were used: G418 (2.5 μg/ml), Hygromycin (0.5 μg/ml), Puromycin (0.05 μg/ml), Blasticidin (2 μg/ml) and Phleomycin (2.5 μg/ml).

### Generation of products required for CRISPR/ Cas9 transfections

The Bar-seq primer design from LeishGEdit ^80,81^ was used to amplify a repair template from the pPOT plasmid. The LeishGEdit primers allow tagging of the wild-type allele or knock-out of the gene. Upstream and downstream, primers contain primer binding sites compatible with pPOT plasmids and 30 nucleotide homology arms for recombination. sgRNA, primers consist of a T7 RNA polymerase promotor (for in vivo transcription of RNA), a 20 nucleotide sgRNA target sequence to introduce the double-strand break at a locus of interest and a 20 nucleotide overlap to the sgRNA backbone sequence allowing the generation of sgRNA templates by PCR ^79,80,82^. We modified the primers in the LeishGEdit protocol to allow us to replace the wild-type gene with a tagged monomorphic or pleomorphic sequence we had previously synthesised. We used the upstream forward primer 1 as per the LeishGEdit protocol and, for the downstream reverse primer, we used the start of primer 2 which binds the GS-linker of the pPOT plasmid and the tail of primer 7 which binds the 3’ UTR of the target gene. All of the replacements were N terminally Ty tagged ^83^ and introduced under 3’ UTR endogenous expression. The 5’ sgRNA and 3’ sgRNA primers were maintained as per the LeishGEdit protocol The LeishGEdit protocol uses the TREU927/4 genome to create primer binding sites. The binding sites of the primer sequences to the endogenous locus were visualised and the binding sites of each were screened for a mutation in the *T. b. brucei* EATRO 1125 genome, based on SNP calls. If necessary, primers were recoded to match the EATRO 1125 sequence. A full list of primers used in this study can be found in Supplementary Data S3.

The repair and sgRNA products were amplified for transfection using Phire Hot Start II DNA Polymerase (Thermo Fisher). The PCR was run for 30 cycles with the following conditions varying for the repair or sgRNA amplification respectively: annealing temperature 65°C and 60°C, 3% DMSO and 0% DMSO and primer concentration 0.5µM and 2 µM.

5µg of each product was used for transfections into the *T. brucei* EATRO 1125 AnTat1.1 J1339 (J1339) pleomorphic cell line ^5^. The J1339 cell line contains the pJ1339 plasmid derived from pJ1173 that carries a puromycin resistance marker, the Tet Repressor, T7 RNA polymerase and Cas9, which is expressed constitutively ^80^. To transfect the parasites, we followed previously described protocol, but modified the Amaxa Nucleofector II using the CD4+ T cells program to Z-001 ^84^.

gDNA from was extracted from clonal transfectants using the DNeasy Blood & Tissue Kit with an RNAse A step (Qiagen). Successful replacements were then validated using the section of primers 1 and 7 which bind gDNA.

### In vitro developmental competence phenotype screen

Replacement clones were seeded at 1×10^5^ cells/ml and grown in either HMI-9 or HMI-9 supplemented with 15% BHI in triplicate. The parasite population density was counted at 0, 24, 48 and 72 hours using the Beckman Coulter Z2 Cell and Particle Counter.

A separate culture was used to generate immunofluorescence images of the gene replacement cell lines. The gene replacement clones were seeded at 1×10^5^ cells/ml and left for 48 hours. Cell cycle status and PAD1 expression were analysed by 4’,6-diamidino-2-phenylindole (DAPI) (100 ng/ml) and an anti-PAD1 antibody ^85^. The cells were washed with vPBS and then resuspended in vPBS and 8% PFA (1:1) and incubated for 10 minutes at room temperature. 2µL of IGEPAL CA-630 (Sigma-Aldrich) 10% in PBS was added and the cells were incubated for a further 10 minutes. The cells were centrifuged at 900g for eight minutes and resuspended in 0.1% glycine in vPBS and incubated for 10 minutes. The cells were then centrifuged at 900g for eight minutes and resuspended in vPBS. Fixed cells were added to a Polysine slide and left to dry. The cells were rehydrated in PBS for five minutes and blocked by incubation of the cells in PBS 2% BSA for one hour. The blocking buffer was removed and anti-PAD1 (1:1000) PBS 0.2% BSA was added to the cells and incubated for one hour. The cells were washed 5x with PBS and then incubated in the Alexa Fluor 568 Goat anti-Rabbit secondary antibody (1:500) (Invitrogen) PBS 0.2% BSA for one hour. 25µL of DAPI (100 ng/ml) in PBS was added to the cells and incubated for five minutes. The cells were then washed five times with PBS. Finally, the slides were mounted with Fluoromount-G Mounting Medium (Invitrogen). All steps were performed at room temperature. The slides were then imaged with a ZEISS Axio Imager 2 using a 40x objective with a Prime BSI Camera. Images were generated with Micro-Manager 2.0 ^86^ and analysed with FIJI ^87^. The number of arrested (1K1N) and dividing (2K1N and 2K2N) ^88^ parasites were counted along with the number of PAD1+ cells.

The motility of FAZ41 replacement cell lines were quantified using 5×10^5^ cells which were centrifuged at 800g for five minutes and resuspended in 40µl of HMI-9. The resuspended cells were then placed onto a slide and a cover slip was mounted using Vaseline. The slides were then incubated for 5 minutes (37°C, 5% CO2). Each slide was visualised at 40x magnification using the Olympus CKX53 and a time-lapse image was generated with a QImaging Retiga-2000R digital camera. 100 images were then taken, one image every 0.25 seconds. The images were imported into FIJI ^87^ and the background was subtracted (rolling ball radius: 10, light background) from the images. The image was then inverted to improve the contrast between the cell and the background. TrackMate ^89^ was used to identify cells using default parameters, unless stated otherwise. LoG detector was used with estimated object diameter: 15 and quality threshold: 0. An initial spot threshold of 0.74 was then applied and the advanced Kalman Tracker to track cell motility (initial search radius: 30, search radius: 30 and max frame gap 20). A summary of each track was then exported for each time lapse.

For analysis of monomorphic cell line responsiveness to BHI, *T. b. brucei* EATRO AnTat1.1, *T. brucei* Lister 497, *T. b. equiperdum* type OVI (OVI isolate) and *T. b. evansi* type A (RoTat 1.2 isolate) were incubated with HMI-9 containing 0%, 1%, 2.5% or 5% BHI for 48 hours. Cell counts were performed using a Neubauer slide (140468, Dominique Dutscher) after 48 h incubation with a Motic AE30 inverted binocular microscope (Motic, Hong Kong, China). Parasite solutions from HMI-9 medium (negative control) and HMI-9 medium supplemented with 5% BHI were used for PAD1 expression analysis. 2×10^6^ trypanosomes were centrifuged at 900g for 5 min. The culture supernatant was removed and the parasite pellet was washed with cold 1X PBS. The washed cells were then fixed with 4% paraformaldehyde (PFA) for 10 min on ice. Cells were resuspended in 130 µl of 1X PBS - 0.1 M glycine and incubated overnight at 4°C. Cells fixed in 4% PFA were then diluted in 1X PBS to a concentration of 1×10^4^ cells/ml. Twenty microliters of 4% PFA-fixed cells were deposited on an 8-well Labtek slide (Dominique Dutscher - 2515364) previously treated with Poly-L-Lysine (P4707 - Sigma Aldrich), and the fixed cells were incubated at room temperature for 1 h. Membrane permeabilization of parasites adhering to the Labtek slide was performed using a PBS - 1X Triton 0.1% solution for 2 min, followed by washing with 1X PBS. Saturation of non-specific sites in Labtek slide wells was performed with PBS 1X - BSA 2% solution for 45 min at 37 °C in a humidity chamber prior to addition of 20 µl anti-PAD1 primary antibody (diluted in PBS 1X - BSA 2%). Wells were washed 5 times with PBS 1X and then incubated with 20 μl of secondary antibody (diluted in PBS 1X - BSA 2%, α-lapin Alexa fluor 488 1:500) for 45 min at 37°C in a humidity chamber before incubation for 10 min with 50 µl of DAPI solution (NucBlue Hoechst Live Cell Stain ReadyProbes, ThermoFischer - R37605). Wells were washed 5 times, then a drop of ProLong® Gold Antifade Reagents (Invitrogen™ P36941) was applied before mounting the slides. Slides were then analyzed on an Axiolab® Zeiss microscope using a 40x objective. Negative controls (no primary antibody and no secondary antibody) and positive controls (*T. brucei* EATRO AnTat1.1 incubated 48 h in HMI-9 medium with 5% BHI) were included for each experiment.

### Monomorph selection and transcriptomic profiling

Pleomorphic *T. brucei* EATRO 1125 AnTat 1.1 90:13 was cloned via serial dilution. In a similar approach to previous studies, the cell line was grown in HMI-9 at 37°C and 5% CO_2_ ^90^ for a total of 72 days ^42^. In this study, we supplemented the HMI-9 with BHI. During the 72-day selection, the percentage of BHI was progressively increased from 2.5% to 15%. The cell density was counted at each passage using a Beckman Z2 Coulter particle counter and size analyser. High cell density was deliberately maintained to select proliferating cells at high parasite population density. Cultures were cryopreserved weekly during the selection period at −80°C in HMI-9 supplemented with 10% glycerol. Before the cultures were screened for their responsiveness to BHI, the cells were washed twice with PBS and grown in HMI-9 for one week to remove any residual BHI from the media.

RNA was extracted from triplicate cultures of the ‘start’ and ‘end’ from the five clones described above. The cells were grown to a density of between 7.5×10^5^ cells/ml and 1×10^6^ cells/ml and RNA was then extracted using the Qiagen RNeasy Mini kit, following the manufacturer’s instructions. RNA was then sequenced with DNBseq (∼4Gb/ sample) by BGI (Hong Kong).

Our analysis was based on a standard workflow for analysing RNAseq data ^91^. Salmon ^92^ was used to index the TREU927/4 reference genome, which was then used to quantify transcript abundance for each sample. Transcript abundance values were imported into R with tximport ^93^. A database was created using GenomicFeatures ^94^. Genes with a count of less than 10 were excluded from the analysis and a VSD transformation was applied to visualise the data as heatmaps and PCA plots. Clones 1 and 3 were removed from the analysis as the replicates did not group via PCA.

DESeq2 ^91^ was used to quantify differentially expressed genes. Differentially expressed genes were identified between the start and end of the selection. For the clonal selection, initially differentially expressed genes were identified by grouping all the clonal libraries. The contrast function was then used to identify differentially expressed genes for each clonal selection using the same model. To compare the similarity in differentially expressed genes between the clonal selections, a Venn diagram was created with Venn (available here https://github.com/dusadrian/venn) to highlight shared differentially expressed genes between the clonal selections. GO enrichment was calculated with TriTrypDB ^95^, including a Benjamini false discovery rate cut-off of >0.05. GO enrichment was visualised with ggplot2 in R.

### Overexpression of ZC3H20 and RBP10

ZC3H20 (Tb927.7.2660) and RBP10 (Tb927.8.2780) were chosen to attempt rescue pleomorphism in the selected monomorphic *T. brucei* cell line. The genes were amplified from pleomorphic *T. brucei* EATRO 1125 genomic DNA. These amplicons were ligated into the TOPO plasmid using T4 ligase (NEB), following the manufacturer’s instructions. The ligated plasmids were then transformed into competent cells and grown overnight. A small-scale plasmid preparation was performed on the culture to isolate the plasmid using the GeneJET Plasmid Miniprep Kit (Thermo Fisher). This plasmid was digested using XbaI and BstXI (5’ tag) or HindIII and SpeI (3’ tag) enzymes (NEB) following the manufacturer’s instructions, and the genes were ligated into the doxycycline-inducible pDex-577y plasmid ^96^. This plasmid integrates into the 177bp repeat sequences of minichromosomes and encodes for phleomycin resistance. ZC3H20-pDEX-577y and RBP10-pDEX-577y were transfected into selected monomorphic clone A7.

The developmental competence of the cell lines transfected with ZC3H20-pDEX-577y and RBP10-pDEX-577y was determined in vitro using BHI-based oligopeptides as above whilst gene overexpression was either uninduced or induced. Doxycycline induction was titrated from 0.2ng/ml to 0.02fg/ml in 1:10 increments.

### In vivo infections

Experiments were carried out according to UK Home Office Animal (Scientific Procedures) Act (1986) under licence number PP2251183. In vivo infections were performed using four female MF1 mice per cell line. Mice were cyclophosphamide treated (25 mg/ml) 24 hours before infection with 10,000 parasites via intraperitoneal injection. Parasitaemia was monitored from 3 days post-infection by tail snip. Blood-smeared slides were air dried at room temperature for five minutes and then fixed via submersion in methanol at −20°C for 10 minutes or stored in methanol at −20°C. PAD1+ and dividing cells were identified using the immunofluorescence protocol described above.

### Statistical analysis

A Wilcoxon two-sample test (Fig. 2f) or a repeated measures ANOVA, including an adjusted post-hoc Bonferroni test (Fig. 2a, Fig. 2c, Fig. 2e, Fig. 3a, Fig. 3b, Fig. 3c and Fig. 4e) was performed using Rstatix ^97^. The data was summarised and checked for normality and outliers before analysis. Full details of statistical results can be found in Supplementary Data S4.

### Data and materials availability

All data and code are available in the main text, the supplementary materials, GitHub (https://github.com/goldrieve/Mechanisms-of-life-cycle-simplification) or NCBI under the BioProject PRJNA1114649.

## Acknowledgments

We thank Dr. Frederik Van den Broeck for facilitating data sharing, Prof. Achim Schnaufer, Prof. Philippe Büscher, Dr. Joanna Young, Prof. Steve Kelly and Dr. Al Ivens for their valuable insights and comments on the work presented here.

## Author contributions

Conceptualization: GRO, KRM

Methodology: GRO, FV, MC, MG, NVR, KRM

Investigation: GRO, FV, MV, KRM

Visualization: GRO, KRM

Funding acquisition: MG, NVR, KRM

Project administration: GRO, FV, KRM

Supervision: MC, KRM

Writing – original draft: GRO, KRM

Writing – review & editing: GRO, FV, MC, LH, MG, NVR, KRM

## Funding

Wellcome Trust (221717/Z/20/Z): Keith R Matthews

Wellcome Trust (220058/Z/19/Z): Guy R Oldrieve, Keith R Matthews

Wellcome Trust (108905/Z/15/Z): Frank Venter, Keith Matthews

MC is supported by a Medical Research Council Career Development Award [MR/W026996/1]. This UK funded award is carried out in the frame of the Global Health EDCTP3 Joint Undertaking.

MG and NVR received financial support from the Bill & Melinda Gates Foundation (grant number OPP1174221) and the Flemish Government EWI SOFI-2018 “Cryptic human and animal reservoirs compromise the sustained elimination of gambiense-human African trypanosomiasis in the Democratic Republic of the Congo”.

MV and LH received financial support from the GIS CENTAURE Recherche Equine and the Regional Council of Normandy

## Supplementary Information is available for this paper

Figs. S1 to S6

Data S1 to S4

Supplementary movies 1-5

## Supplementary Materials

**Fig. S1.**
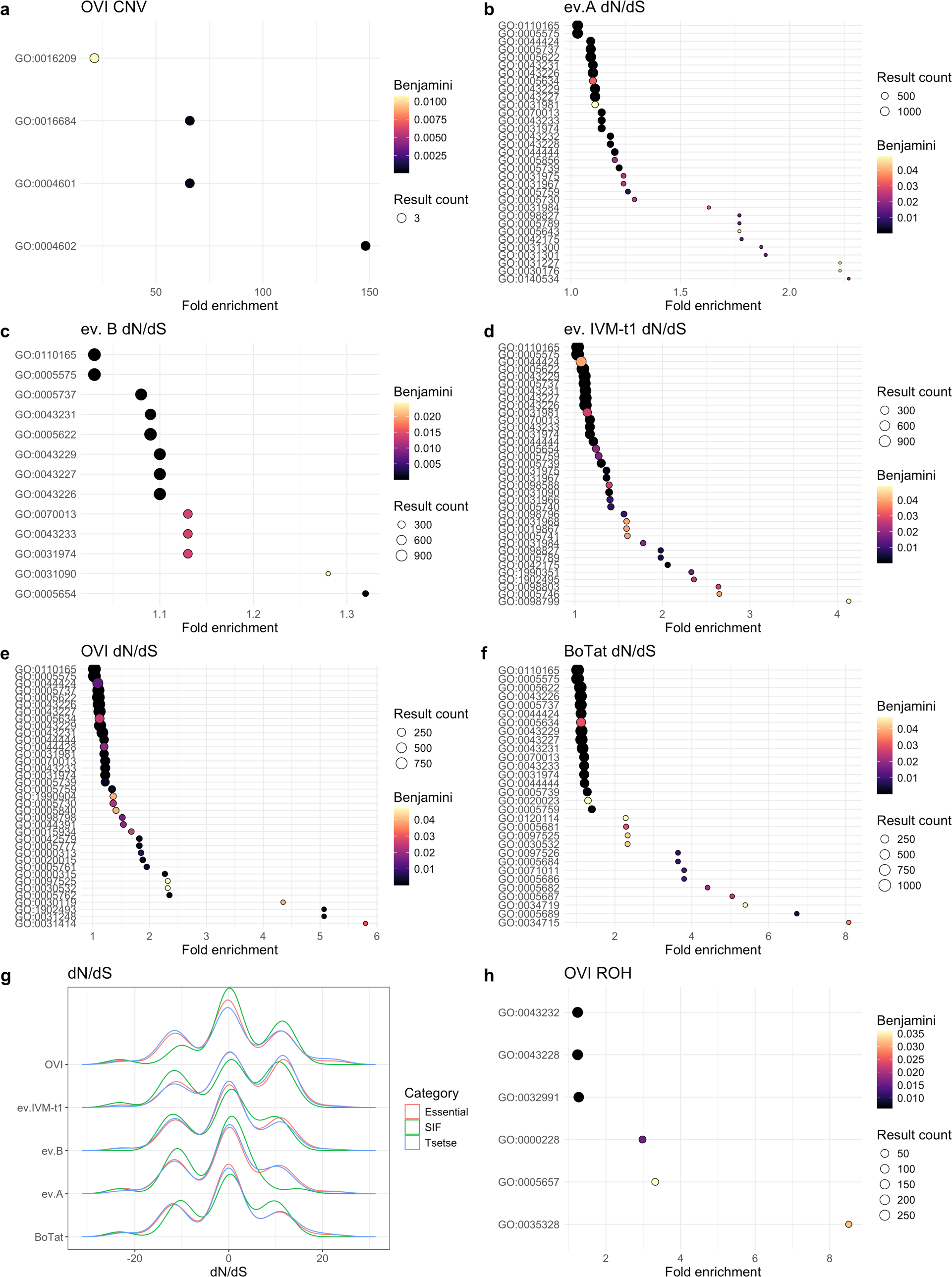
Gene ontology enrichment. **(A)** *T. b. equiperdum* type OVI CNV ‘glutathione peroxidase activity’ (GO:0004602), ‘peroxidase activity’ (GO:0004601), ‘oxidoreductase activity acting on peroxide as acceptor’ (GO:0016684) and ‘antioxidant activity’ (GO:0016209). **(B)** *T. b. evansi* type A dN/dS; mitochondrion (GO:0005739) and mitochondrial matrix (GO:0005759), **(C)** *T. b. evansi* type B dN/dS; ‘membrane-bounded organelle’ (GO:0043227), ‘intracellular membrane-bounded organelle’ (GO:0043231), ‘membrane-enclosed lumen’ (GO:0031974) and ‘organelle membrane’ (GO:0031090**. (D)** *T. b. evansi* type IVM-t1 dN/dS; mitochondrion (GO:0005739), mitochondrial envelope (GO:0005740), mitochondrial membrane (GO:0031966), mitochondrial matrix (GO:0005759), mitochondrial outer membrane (GO:0005741), mitochondrial respirasome (GO:0005746), outer mitochondrial membrane protein complex (GO:0098799). **(E)** *T. b. equiperdum* type OVI dN/dS; mitochondrial large ribosomal subunit (GO:0005762), mitochondrion (GO:0005739), mitochondrial matrix (GO:0005759), mitochondrial ribosome (GO:0005761), mitochondrial protein-containing complex (GO:0098798). **(F)** *T. b. equiperdum* type BoTat dN/dS; mitochondrion (GO:0005739), mitochondrial matrix (GO:0005759). **(G)** dN/dS ratio variation for genes associated with the tsetse fly stage, QS pathway and essential genes, defined by phenotypes in each of the four RITseq libraries; monomorphic *T. brucei* have clade-specific variation in the efficacy of selection. **(H)** *T. b. equiperdum* type OVI ROH ‘transcriptionally silent chromatin’ (GO:0035328), ‘replication fork’ (GO:0005657) and ‘non-membrane bound organelle’ (GO:0043228).

**Fig. S2.**
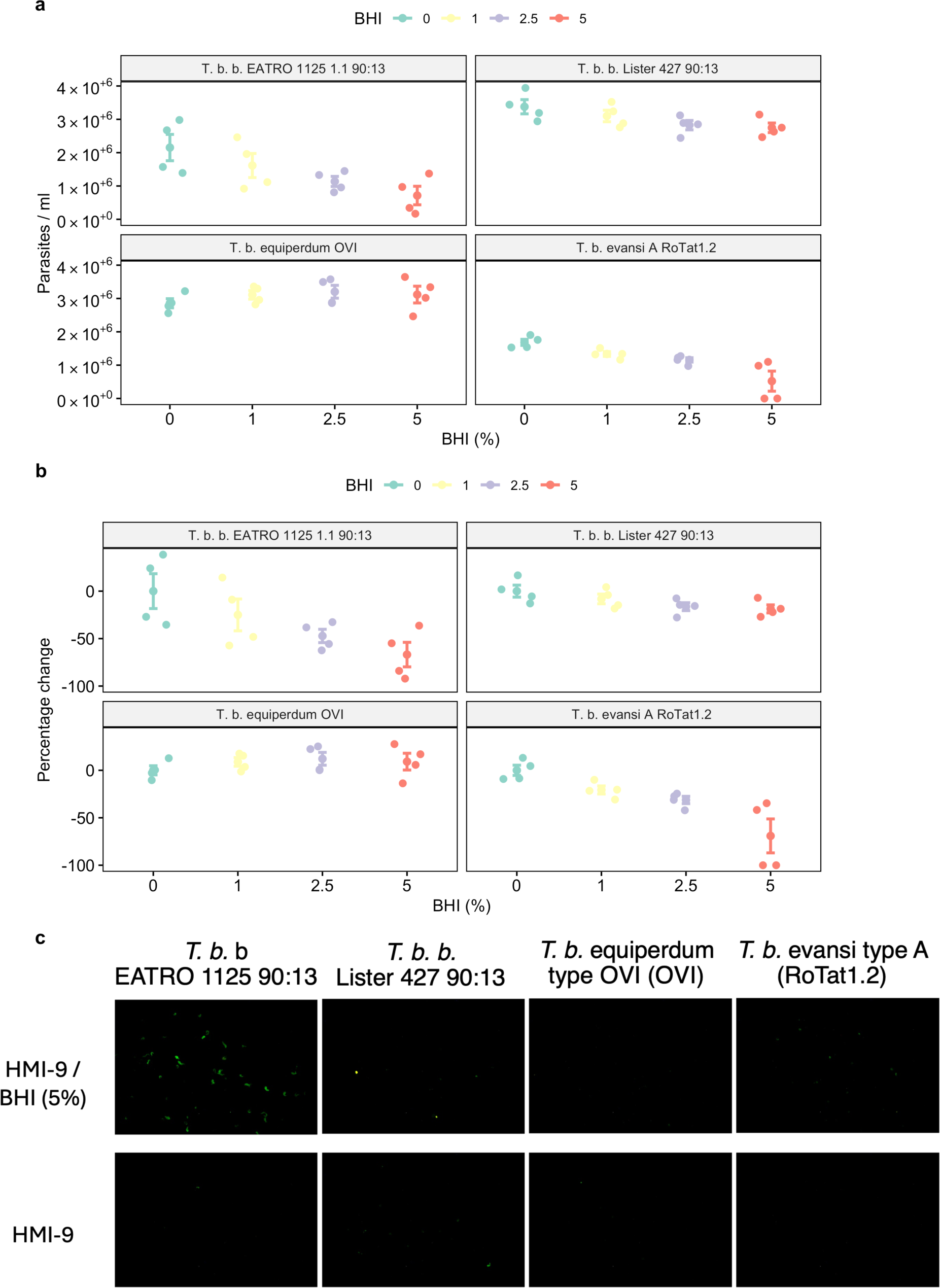
Growth of *T. brucei* strains exposed to BHI supplemented media for 48 hours. Pleomorphic *T. b. brucei* EATRO 1124 90:13 and monomorphic *T. b.* Lister 427, *T. b. equiperdum* type OVI (OVI isolate) and *T. b. evansi* type A (Rotat1.2 isolate) were exposed to BHI at either 0, 1, 2.5 or 5 percent. The data was compared as **(A)** parasite density or **(B)** percentage change, calculated as the density of each replicate at each BHI concentration compared to the mean density at 0% BHI for the corresponding strain. **(C)** Representative PAD1 images of each cell line grown in 0% and 5% BHI.

**Fig. S3.**
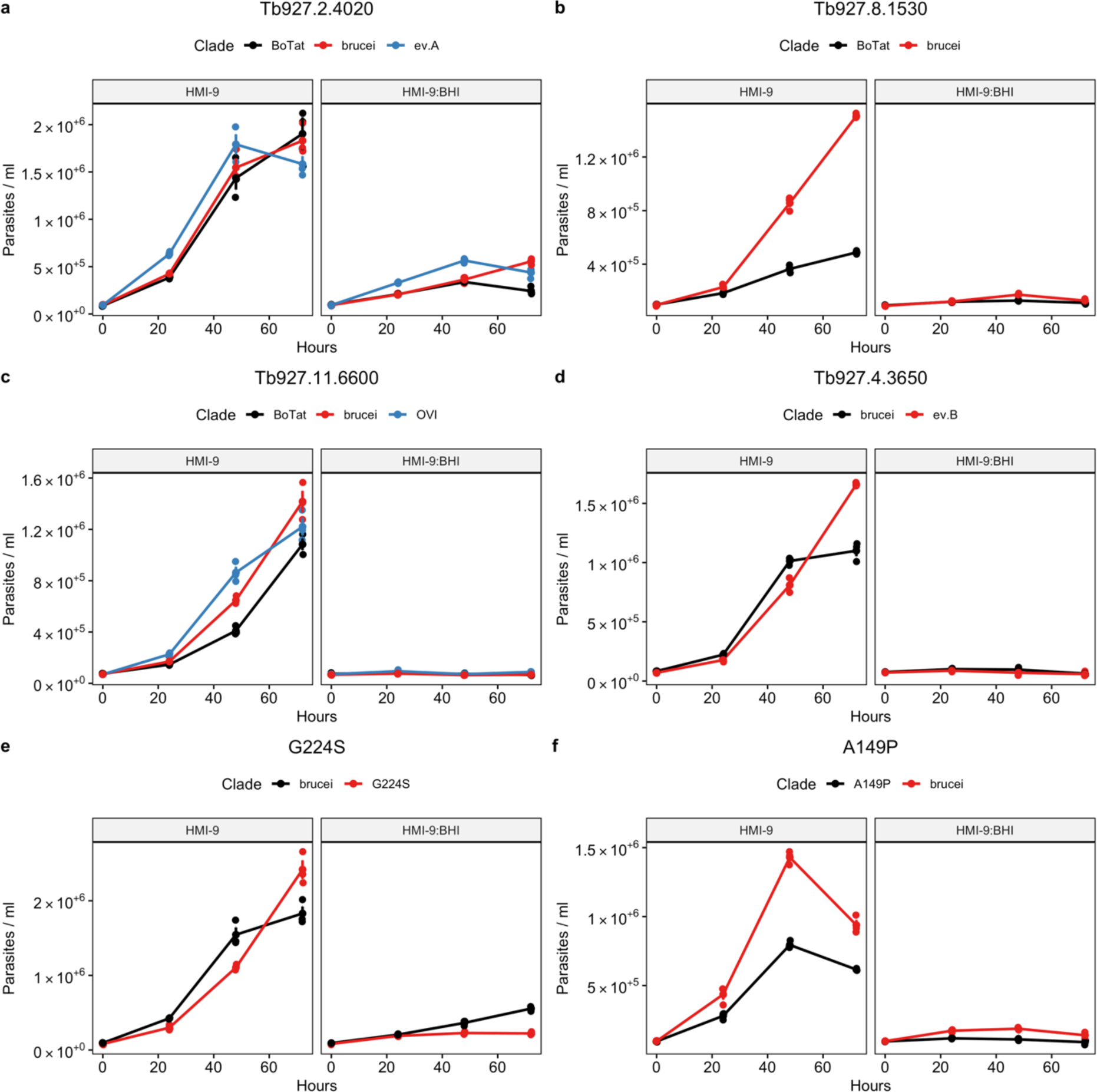
Growth of pleomorphic *T. brucei* EATRO 1125 AnTat1.1 subjected to replacement of endogenous gene targets. **(A)** Tb927.2.4020 APPBP1, **(B)** Tb927.8.1530 - Golgi pH regulator (GPR89), **(C)** Tb927.11.6600 – Hyp1, **(D)** Tb927.4.3650 – Protein phosphatase 1 (PP1), **(E)** G224S, **(F)** A149P. The replacement cell lines were grown in HMI-9, or HMI-9 supplemented with an in vitro mimic of the QS signal, oligopeptide broth BHI (15%).

**Fig. S4.**
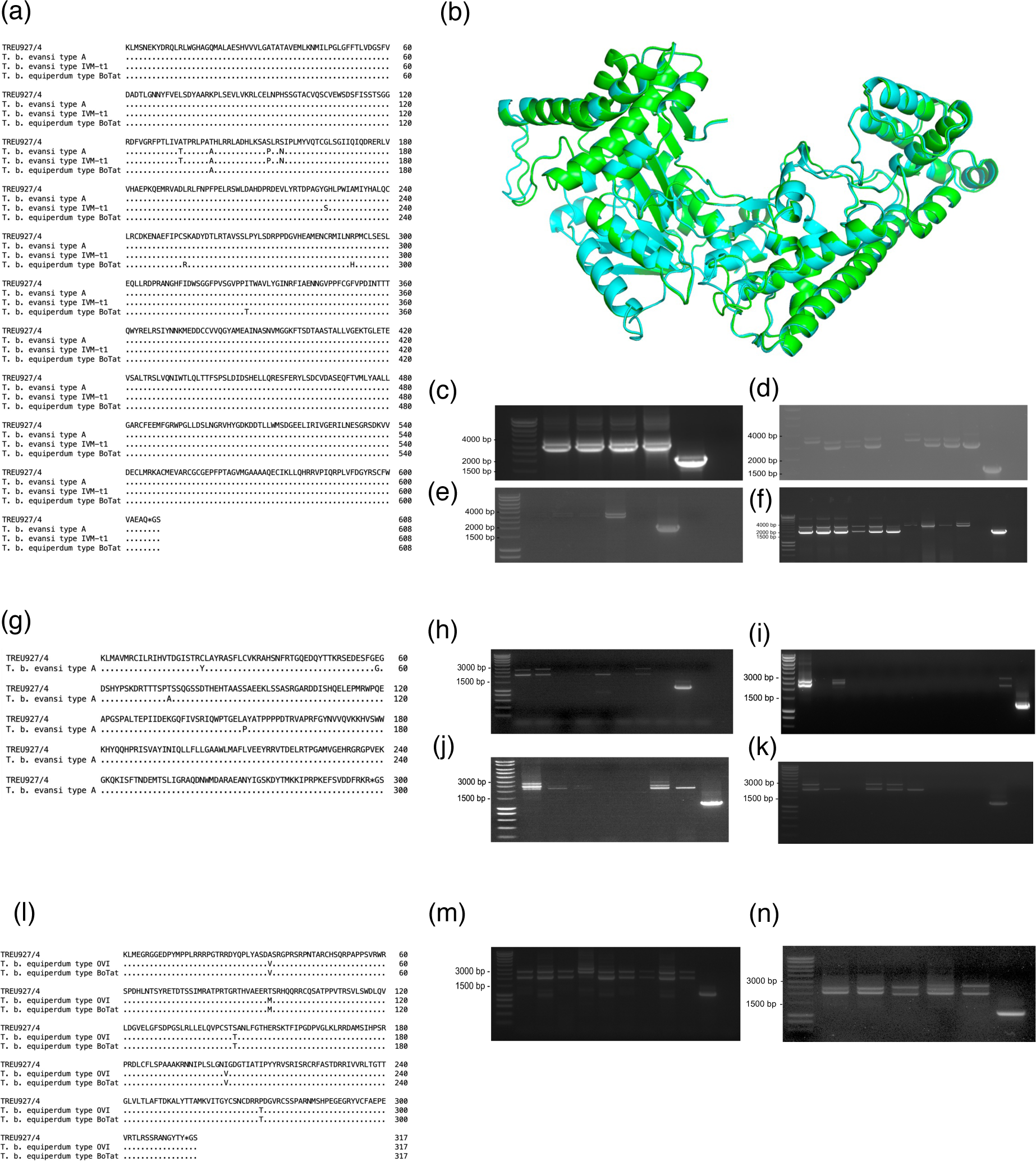
Confirmation of replacement of endogenous gene targets in pleomorphic *T. brucei* EATRO 1125 AnTat1.1. Tb927.2.4020 **(A)** gene alignment **(B)** *T. b. brucei* (blue) and *T. b. evansi* type IVM-t1 (green) protein structure alignment (RMSD = 0.210). **(C-F)** PCR confirmation of endogenous allele replacement. **(C)** Replacement of wild-type APPBP1 (Tb927.2.4020) with the *T. b. equiperdum* type BoTat sequence (clone 4, lane 1*), T. b. evansi* type A sequence (clone 1, lane 2), *T. b. evansi* type IVM-t1 sequence (clone 3, lane 3), *T. b. brucei* sequence (clone 1, lane 4) and control J1339 (lane 5). **(D)** Heterozygous phleomycin add-back (clones 1-4, lanes 1-4), Heterozygous G418 add-back (clones 1-4, lanes 6-9) and control J1339 (lane 10). **(E)** Homozygous phleomycin+G418 add-back (clones 2-3, lanes 1-2), homozygous G418+phleomycin add-back (clones 2-3, lanes 3-4) and control J1339 (lane 6). **(F)** G224S heterozygous replacement (clones 1-6, lanes 1-6) and homozygous replacements (clones 1-4, lanes 7-10) and control J1339 (lane 12). Expected sizes (bp): Wild type Tb927.2.4020 = 1,992, pPOT-Hygromycin-Tb927.2.4020 = 3,425, pPOT-Blasticidin-Tb927.2.4020 = 2,996, pPOT-Phleomycin-Tb927.2.4020 = 2,221, pPOT-Neomycin-Tb927.2.4020 = 3,194. Tb927.5.2580 **(G)** gene alignment **(H-K)** PCR confirmation of endogenous allele replacement. **(H)** Replacement of wild-type Tb927.5.2580 with *T. b. evansi* type A sequence (clones 1-3, lanes 1-3) and the *T. b. brucei* sequence (lanes 4-8, clones 1-5) and control J1339 (lane 10). **(I)** A149P sequence (clones 1-7, lanes 1-7) and control J1339 (lane 10). **(J)** heterozygous add-back (clones 1-11) and control J1339 (lane 12). **(K)** Homozygous add-back (clones 1-8) and control J1339 (lane 9). Expected sizes (bp): Wild type Tb927.5.2580 = 1,134, pPOT-Hygromycin-Tb927.5.2580 = 2,337, pPOT-Blasticidin-Tb927.5.2580 = 1,908, pPOT-Phleomycin-Tb927.5.2580 = 1,893, pPOT-Neomycin-Tb927.5.2580 = 2,106. Tb927.11.3400 **(L)** gene alignment **(M-N)** PCR confirmation of endogenous allele replacement. **(M)** Replacement of wild-type Tb927.11.3400 *T. b. equiperdum* type BoTat (clones 1-3, lanes 1-3) sequence, *T. b. brucei* (clones 1-3, lanes 4-6) sequence and *T. b. equiperdum* type OVI (clones 1-3, lanes 7-9) sequence and control J1339 (lane 10). **(N)** *T. b. equiperdum* type OVI homozygous add-back (clone 1-2, lanes 1-2), *T. b. equiperdum* type BoTat homozygous add-back (clone 1-2, lanes 3-4), *T. b. brucei* initial replacement control (lane 5) and control J1339 (lane 6). Expected sizes (bp): Wild type Tb927.11.3400 = 1,110, pPOT-Hygromycin-Tb927.11.3400 = 2,552 and pPOT-Blasticidin-Tb927.11.3400 = 2,123, pPOT-Phleomycin-Tb927.11.3400 = 2,108, pPOT-Neomycin-Tb927.11.3400 = 2,321.

**Figure S5.**
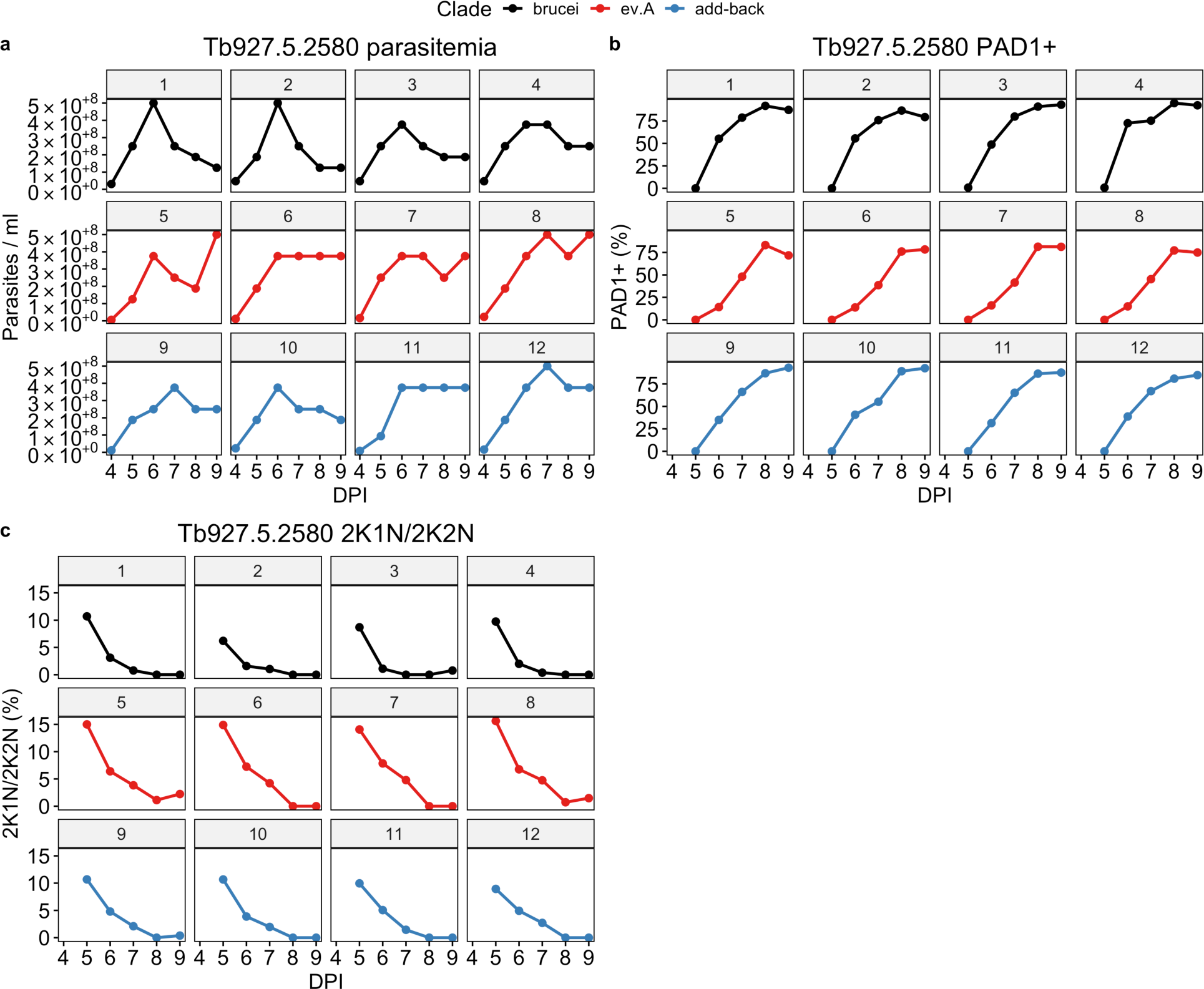
Pleomorphic *T. b. brucei* expressing the monomorphic *T. b. evansi* type A Tb927.5.2580 sequence delays developmental progression in vivo. **(A)** In vivo growth of pleomorphic *T. b. brucei* expressing the monomorphic *T. b. evansi* type A Tb927.5.2580 sequence or the pleomorphic *T. b. brucei* sequence, **(B)** cell cycle stage and **(C)** percentage of PAD1+ cells as assessed by immunofluorescence. Each cell line was used to infect four mice, represented by each panel.

**Figure S6.**
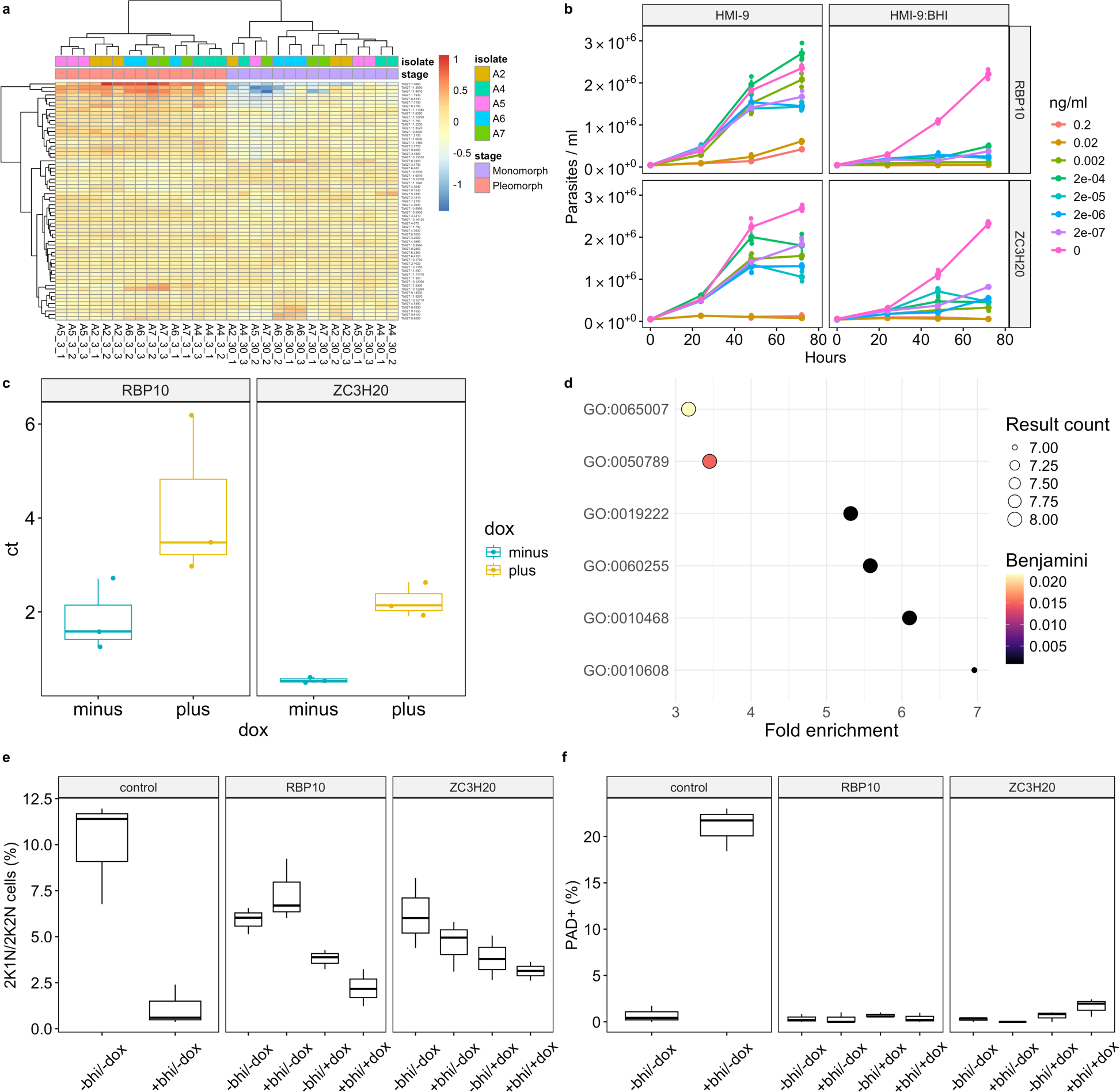
Selected monomorphs display reduced developmental progression, which is reversible by inducible expression of key gene regulators. **(A)** Heatmap of SIF genes. Normalisation was achieved by calculating the deviation of expression from each gene in a specific sample from the mean expression of that gene across all samples. Blocks of genes and samples which covary are clustered, indicated by branches. The split between each stage of the selection was not specified. **(B)** Growth of clone A7 in HMI-9 or HMI-9: BHI induced to overexpress RBP10 or ZC3H20 at titrated concentrations of doxycycline **(C)** Quantification was calculated via RT-qPCR using ZC3H20 or RBP10 specific primers and cDNA generated from clone A7 induced to overexpress either gene. The expression of the gene of interest was normalised to ZFP3 and compared between uninduced (0ng/ml) and induced (0.002ng/ml) cultures grown in triplicate (2^-(ΔΔCT))**. (D)** GO molecular function enrichment from 23 commonly differentially expressed transcripts included significant enrichment of ‘regulation of gene expression’ (GO:0010468), ‘posttranscriptional regulation of gene expression’ (GO:0010608) and ‘regulation of macromolecule metabolic processes’ (GO:0060255). **(E & F)** percentage of the population replicating and PAD1+% from the monomorphic clone A7 induced with doxycycline to express transgenic RBP10 and ZC3H20 or not and grown in HMI-9 or HMI-9 supplemented with 15% BHI.

## Supplementary data descriptions

**Data S1:**

Database of trypanosome isolates, sequencing metrics, region of origin and sequencing technology used to derive the genomic datasets in this study.

**Data S2:**

Clade-specific variants were prioritised to create a target list of genes to validate their role in monomorphism. Initially, pseudogenes, VSGs and genes not on the megabase chromosomes 1-11 were removed. These genes were further filtered to create two target categories using the following criteria:

Category 1 (Tab 1) Genes which have a clade specific high-impact or moderate-impact mutation in a monomorphic clade in a known QS pathway gene. The gene must also display no high impact mutations in any pleomorphic isolates.

Category 2 (Tab 2). Genes which have a clade specific high-impact or moderate-impact mutation in a monomorphic clade along with a dN/dS ratio <1 in pleomorphic isolates and >1 in any monomorphic clade. The gene must also have a smaller log fold change in D3, D6 and PF than in the DIF category whilst displaying a log fold change in the DIF category greater than −1.5. The gene must also display no high impact mutations in any pleomorphic isolates.

(Tab 3) genes with a dN/dS ratio >1 in all monomorphic clades and <=1 in the pleomorphic background.

**Data S3:**

Oligonucleotide primer and antibody resources used in this study.

**Data S4:**

Statistical reporting for all Figures.

**Supplementary movies 1-5:**

Cell motility video tracks for *T. brucei* expressing Tb927.11.3400 derived from BoTat or OVI isolates, or with add back of the wild type sequence.

